# Comparative analysis of microbial communities of soils under contrasting microclimates

**DOI:** 10.1101/2024.11.27.625671

**Authors:** T Päll, E Zagal-Venegas, T Pavlicek, E Nevo, S Timmusk

**Author notes:** **Correspondence** Salme Timmusk, Dept. of Forest Mycology and Pathology, Uppsala BioCenter, Box 7026 SE-75007.

## Abstract

Understanding how microbiomes influence the life cycle and fitness of crops, and how global change drivers disrupt this network, is pivotal for an understanding of the crop as a holobiont, and of how to provide solutions for Nordic agricultural crop resilience under climate change.

Despite decades of use of plant growth-promoting rhizobacteria (PGPR), there is an intrinsic problem with their applications, as it has become evident that their functionality and performance rely on interactions with the environment and with other microorganisms. The synthetic crop-promoting rhizobacterial community strains are being outcompeted by native communities, or their colonisation and active principles are being reduced to ineffective levels. This is the result of the communities being selected on taxonomic criteria rather than qualitative analysis of the microbiome-associated plant phenotypes. In this context there in an urgent need for an approach studying the microbial community and plant complementarity traits from indigenous communities. Here we report the pattern of bacterial distributions at the Evolution Canyon (EC) in Israel to gain insight into microbiomes exposed to contrasting microclimates at the North Facing Slope (NFS) and South Facing Slope (SFS) sun and shade areas using high-throughput sequencing. While the NFS and SFS shaded areas bacterial distribution didn’t differ, our results show significant differences between the NFS and the SFS sunny areas. The families *Geodermatophilaceae, Beijerinckiaceae*, and *Pseudonocardiaceae* are dominant in the NFS sun area, and the families *Rubrobacteriaceae*, unclassified *Solirubrobacterales* bacterium 67-14, unclassified *Actinobacteriota*, class *Gaiellales* dominate at the SFS sun area. Likewise, both Shannon and inverse Simpson’s diversity indices are higher at the NFS sun area compared to the NFS shaded area. There was no substantial difference between diversity indices in SFS sun and shaded area.

Our results advance our understanding of the bacterial distributions at what is in effect a natural laboratory of ecosystems that probably evolved 5–7 million years ago. The data are an important step towards using transcriptomics, metabolomic profiles and selective plating for figuring out key strains and the supporter strains that strengthen the ecological functions of the key strains. Collectively, this will enable us to assemble redundant and stable synthetic PGPR communities consisting of key and supporter strains for promoting plant health and stress tolerance under changing climates.

## 1. Introduction

Climate change, stagnating crop yields and the increased demand for food pose challenges to future agriculture, and it is of great importance to assess the potential of agricultural management practices to mitigate the effects of climate change on crop productivity.

Healthy soils with functional microbiomes are crucial for any sustainable production system. Soils harbour a substantial fraction of the world’s biodiversity, contributing to many critical ecosystem functions, and are heterogeneous environments with various dynamic parameters that can affect microbial growth and survival. While past efforts have focused on weather, microbial distributions have geographic or spatial patterns, currently, available sequencing methods can lead to further understanding of spatial, environmental, taxonomic, and functional patterns important for microbes across ecosystems and biomes. The Earth is covered by ∼ 10^30^ microorganisms from roughly 10^12^ species that have resulted from ∼4 billion years of evolution. Microorganisms represent the largest fraction of the global biomass, and bacteria are usually the dominant microorganisms [1, 2]. The history of microbial commercial application goes back more than 100 years when plant growth-promoting rhizobacteria (PGPR) were used to influence biotic fitness and fight agricultural pathogens. The paradigm change occurred in 1999 when it was discovered that in addition to fighting biotic stress factors, PGPR can influence plant abiotic stress conditions by enhancing desiccation tolerance [3]. Huge numbers of scientific and commercial articles and patents have been published regarding how to apply PGPR both as a biocontrol measure and as a general growth promoter in agricultural settings. Despite decades of use, there is an intrinsic problem with PGPR applications, as the products have limited persistence under field conditions [4, 5], and the products in the natural environment often don’t provide the same benefits as under controlled conditions [4-12]. PGPR strains are being outcompeted by native communities, or their colonization and active principles are reduced to ineffective levels [13, 14]. This is the result of crop plant microbiomes being selected on taxonomic criteria rather than qualitative analysis of the microbiome-associated plant phenotypes [9].

Against this background, there is an urgent need for a “back to the roots” approach, studying the microbial community and plant complementarity traits from indigenous communities [1, 5, 9, 10, 15-17]. How do bacteria distribute and speciate in natural environments? Most insight has come from the theoretical and computational experimental laboratory systems in studies of microevolution. These studies, however, do not take into account the diversity of modes of how bacteria can evolve under the complexity of real-world natural settings, including integration with the host organisms. Therefore, for a comprehensive understanding of bacterial spatial distribution and potential functions, it is essential to study natural populations [1]. Insights into the complex interactions of traditional, wild and harsh ecosystems will help improve our understanding of the evolutionary and ecological diversification that controls and improves plant fitness, while these ecosystems can be seen as reservoirs of genetic potential that may be mined [1]. One such well-studied native environment is the natural laboratory called the Evolution Canyon (EC) found in northern Israel [17-19]. The ‘African’ or south-facing slopes (AS or SFS) in canyons north of the equator receive higher solar radiation than the adjacent ‘European’ or north-facing slopes (ES or NFS). The difference in solar radiation causes higher maximal and average temperatures and evapotranspiration on the more stressful ‘African’ slope. Earlier, we studied the bacterial distribution on the opposing slopes. We characterized the microbial spore-forming isolates belonging to five taxa based on metabolic properties: ACC deaminase content, biofilm-forming properties, halotolerance and phosphorus solubilization ability [17]. Our results indicated significantly higher levels of ACC utilization, biofilm formation, phosphorus solubilization and moderately halophilic bacteria from rhizosphere samples from the SFS slope in comparison to those from the NFS slope. However, the sun versus shaded area did not make much difference in any of the measured characteristics [17]. The dramatic difference in the isolates’ metabolic properties raised interest in characterizing the microbial inter- and intraspecific profiles.

In the current study, we used high-throughput sequencing: 16S rRNA analysis of V3-V4 regions to study community structure and microbial diversity in the SFS NFS sun and shaded areas of the wild barley rhizospheres.

## 2. Material and Methods

### 2.1. The Evolution Canyon

Evolution Canyon (EC) is in northern Israel [1, 17-19].The south-facing slopes (SFS) in canyons north of the equator receive higher solar radiation than the adjacent north-facing slopes (NFS). The difference in solar radiation causes higher maximal and average temperatures and evapotranspiration on the more stressful ‘African’ slope. This results in dramatically diverging physical and biotic interlopes, probably originating several million years ago after mountain uplifts. These canyons are remarkable natural evolutionary laboratories. While microclimate remains the major interslope divergent factor, geology, soils, and topography are similar on opposite slopes (50–100 m apart at the bottom). Thus far, to unravel the link between environmental stress and adapting genome evolution, the intraspecific interslope divergence has been compared in 2,500 species across various life forms, from prokaryotes to eukaryotic lower and higher plants, fungi, and animals. The special features of the ecology facilitate drawing up models of biodiversity and genome evolution from which follow testable predictions [17-19].

### 2.2. Rhizosphere sampling and sample preparation

Wild barley rhizospheres were sampled as described previously [17]. Five wild barley plants were collected from each SFS (SFS1, SFS2 and SFS3) and NFS (NFS5, NFS6 and NFS7) sun and shaded stations at EC, two weeks before maturation. The SFS has stations 1, 2, 3 and at 120, 90, and 60 meters above sea level. Three stations on the NFS (5, 6 and 7) are 60, 90, and 120 meters above sea level, respectively. The plant roots were slightly shaken to remove loosely attached soil and to collect soil intimately linked to the plant root. The rhizosphere material (i.e. lateral roots with the attached soil) was placed in new plastic bags, transferred to the laboratory, and then stored at +4°C until they were processed the next day. Plant rhizosphere material (1 g) was homogenized, and DNA was extracted.

### 2.3. DNA extraction and PCR amplification

DNA was extracted from the rhizosphere material at the SFS (SFS1, SFS2 and SFS3) and NFS (NFS5, NFS6 and NFS7) sun and shaded stations. Microbial DNA was obtained from 200 mg of rhizosphere material using the Nucleo Spin® Soil kit (Macherey-Nagel, Germany). Quantitative determination of the concentration (ng/μl) and purity of DNA (A260/280) was performed spectrophotometrically using a NanoDrop Lite Spectrophotometer (Thermo Fisher Scientific). DNA concentration was determined fluorometrically using Qubit 2.0 (Invitrogen). The quality of the DNA was checked via 1% agarose gel electrophoresis. DNA samples were stored at -20°C for further analysis. The routine described earlier for preparing samples for sequencing the variable V3 and V4 regions of the 16S rRNA gene was used [2]. Prokaryotic 16S rRNA gene fragments were amplified by a two-step PCR procedure where the first step consisted of 2.5 ng extracted DNA, 2Х Phusion PCR Mastermix (Thermo Scientific, Waltham, MA, US) and 10 μM of the primers pro341F/pro805R in 15 μl reactions. Two independent PCRs were run, and the PCR products were then pooled. A single 30 μl reaction was performed for the second PCR, using 2 μM of primers with Nextera adaptor and index sequences, and 3 μl of the pooled PCR product from the first PCR. The final PCR products were purified using an E.Z.N.A.® Cycle-Pure Kit (Omega Bio-tek, Georgia) following the manufacturer’s instructions. Sequencing was performed on an Illumina MiSeq instrument using the 2,250 bp chemistry.

### 2.4. Processing of the reads, clustering and taxonomic identification

16S rRNA analysis of MiSeq sequencing data was performed using Mothur v1.48.0, essentially as described in Mothur MiSeq SOP. Paired-end reads were merged with mothur “make.contigs” command using default parameters and pro341F/pro805R oligo sequences. The mean contig length was 414 bases. Merged reads were trimmed using the “screen.seqs” command with “maxambig=0, minlength=402, maxlength=428, maxhomop=8 “parameters. Duplicate sequences were removed using the “unique.seqs” command and aligned to reference alignment. The reference alignment was generated from the Silva reference release v138.1 (https://mothur.s3.us-east-2.amazonaws.com/wiki/silva.nr_v138_1.tgz) using the “pcr.seqs” command with “start=6388, end=25316, keepdots=F” parameters. After alignment to reference, poorly aligned sequences and long homopolymer stretches were removed using the “screen.seqs” command with parameters “count=current, start=40, end=17052”. Overhangs on either side of the v3v4 region were removed using the “filter.seqs” command with parameters “vertical=T, trump=.”. Almost identical sequences were merged using the “pre.cluster” command with the “diffs=3” parameter. Chimeric sequences were identified and removed from further analysis using the “chimera.vsearch” command with the “dereplicate=t” parameter. Retained sequences were classified taxonomically using the “classify.seqs” command using previously generated reference alignment and Silva nr_v138.1 taxonomy. Sequences classified as belonging to taxa “Chloroplast-Mitochondria-unknown-Archaea-Eukaryota” were removed from further analysis using the “remove.lineage” command. Sequences were clustered to OTUs using the distance matrix, generated using the “dist.seqs” command, and the “cluster” command was used to assign sequences to OTUs, both commands were run with the “cutoff=0.03” parameter. OTU read counts were obtained using the “make.shared” command and the “label=0.03” parameter. Consensus taxonomy was assigned to OTUs using the “classify.otu” command and the “label=0.03” parameter. For the downstream analysis, the Mothur final consensus “taxonomy” and the “shared” file with the read counts were used to generate a biom v1.0 file with the R phyloseq v1.48.0 package. The Mothur batch file used to analyse the dataset is shown in the SI 1 file

### 2.5. Data analysis and statistics

Data analysis and figures were done in R v4.4.1 using tidyverse v2.0.0, here v1.0.1, patchwork v1.3.0, ggpubr v0.6.0, ggdist v3.3.2, cowplot v1.1.3, ggnewscale v0.5.0, phyloseq v1.48.0, microViz v0.12.4, microbiome v1.26.0, vegan v2.6-6.1, tidybayes v3.0.6, modelr v0.1.11 R packages. For Bayesian modelling, we used the R libraries rstan v2.21.3 (Stan Development Team 2020) and brms v2.16.1. Models were specified using extended R lme4 formula syntax as implemented in the R brms package. We used weak priors to fit models. We ran a minimum of 2000 iterations and four chains to fit models. Hypotheses were tested using the alpha = 0.05 level, specifying the Bayesian confidence interval (credible interval), containing 1 - alpha = 0.95 (95%) of the posterior values, to determine the presence of an effect.

## 3. Results

We sequenced the procaryotic 16S rRNA gene fragment PCR amplified with oligos encompassing V3-V4 hypervariable regions from 40 samples collected from the SFS, NFS sun, and shaded stations at the EC. The processed raw experiment contained 1,494,590 reads, with a median of 35,655 reads per sample (range 4,560 – 179,293). The number of OTUs in the raw experiment, clustered using a 97% identity threshold, was 56,931, from these, 41,373 (72.7%) were singletons. After removing noninformative low-abundance OTUs using a 0.25% abundance threshold [20], the experiment contained 1,204,499 reads, with a median of 28,604 reads per sample (range 3,016 – 152,345). After low abundance OTU removal, 441 OTUs, including zero singletons, remained for further analysis. Analysis of OTU prevalence showed that 98% (432) OTUs were present in all four stations.

### 3.1. The wild barley rhizosphere microbial communities

Taxonomic classification of the OTUs showed that a total of 13 bacterial phyla and 97 families were identified within our samples and, on average, the three most abundant phyla in the communities were Actinobacteriota 58% (SD = 8%), Firmicutes 19% (8%), Proteobacteria 15% (5%), together accounting for approximately 90% of reads, followed by Chloroflexi 3% (1%), Myxococcota 1% (1%), Acidobacteriota 1% (1%) (<u>Fig. 1A</u> and B). The most abundant families within our samples were Bacillaceae from phylum Firmicutes, accounting for 18% (8%) of reads, Rubrobacteriaceae 12% (5%), unclassified Solirubrobacterales bacterium 67-14 8% (2%), Geodermatophilaceae 5% (3%) from phylum Actinobacteria, and family Beijerinckiaceae 5% (2%) from phylum Proteobacteria (<u>Fig. 1C and D</u>).

**Fig. 1:**
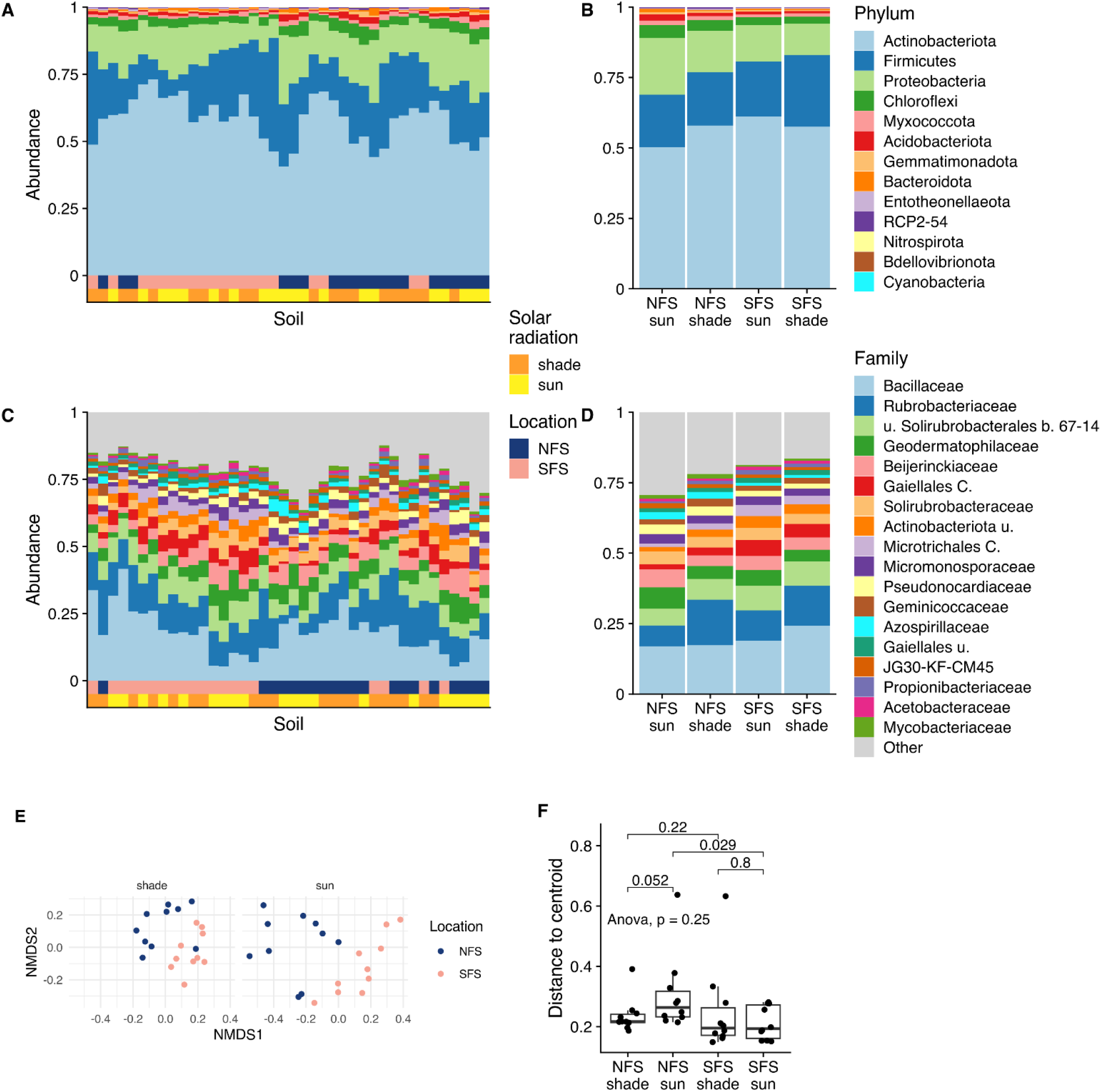
The wild barley rhizosphere microbial communities. Relative abundance of bacterial phyla and families in samples, ordered by Bray-Curtis similarity (A, C). Relative abundance of phyla and families in the SFS, NFS, sun and shaded stations (B, D). A total of 97 families were identified; the top 18 families are shown, and the remaining were merged into the category ‘Other’. E) NMDS ordination of communities based on Bray-Curtis distances of OTU compositional abundances. Points denote individual samples. F) Distances to centroids, calculated from Bray-Curtis distances. Pairwise comparison p-values are from the Wilcoxon signed-rank test; pairwise p-values were adjusted for multiple comparisons using the ‘holm’ method. C., class, b. bacterium, u., unclassified.

Ordination analysis using Bray-Curtis distances of compositional abundances indicated the presence of distinct bacterial communities in the wild barley rhizospheres originating from NFS and SFS stations (Fig. 1 A, C, E). Analysis of the sample pairwise distances within the four stations indicated that NFS sun stations have a higher beta diversity than SFS sun stations (Fig. 1F).

### 3.2. Differential abundance of bacterial taxa in NFS SFS rhizosphere communities

Modelling indicates that the abundance of reads from OTUs assigned to the phylum Actinobacteriota was higher in SFS sun stations compared to the NFS sun stations (Post. Prob. (PP) = 0.989; Fig. 2A and B). In contrast, the OTUs assigned to the phylum Proteobacteria were represented in higher abundance in NFS sun stations compared to the SFS sun stations (PP = 0.992; Fig. 2A and B). There was no substantial difference in the abundance of OTUs assigned to the phyla Actinobacteriota and Proteobacteria in the shaded NFS and SFS stations. The reads of the OTUs assigned to the phylum Chloroflexi were overrepresented in both NFS sun and shade stations compared to SFS sun and shade stations, respectively (PP = 0.974 and 0.97, respectively). Also, OTUs assigned to the phylum Acidobacteriota were slightly more abundant in the NFS sun stations than in the SFS sun stations (PP = 0.966). OTUs assigned to the top three bacterial families – Bacillaceae, Rubrobacteriaceae, and unclassified Solirubrobacterales bacterium 67-14 – were all more abundant in the shaded compared to the sun stations (Fig. 2C). Differential abundance analysis at the bacterial family level indicated bigger and more robust differences in abundance between NFS SFS sun stations than in shaded stations (e.g. more near one PPs for contrasts in sun stations compared to the shaded stations; Fig. 2D). The abundance of the OTUs assigned at the family level to Rubrobacteriaceae (PP = 0.997), unclassified Solirubrobacterales bacterium 67-14 (PP = 0.996), unclassified Actinobacteriota (PP = 1), class Microtrichales (PP = 1), class Gaiellales (PP = 1), and Propionibacteriaceae (PP = 0.998) were more abundant in the SFS sun compared to the NFS sun stations (Fig. 2D). The families unclassified Solirubrobacterales bacterium 67-14 (PP = 0.987), class Gaiellales (PP = 0.999), unclassified Actinobacteriota (P = 0.997), and class Microtrichales (P = 0.994) were also more abundant in the SFS shaded compared to the NFS shaded stations. The families Geodermatophilaceae (PP = 0.992), Beijerinckiaceae (PP = 0.991), Pseudonocardiaceae (PP = 0.999), Azospirillaceae (PP = 0.999), Micromonosporaceae (PP = 0.961), Mycobacteriaceae (P = 0.997), and JG30-KF-CM45 (P = 0.994) were more abundant in the NFS sun compared to the SFS sun stations. From these, the families Pseudonocardiaceae (P = 0.994), Mycobacteriaceae (P = 0.999), Azospirillaceae (P = 0.996), and JG30-KF-CM45 (P = 0.97) were also more abundant in NFS shaded compared to SFS shaded stations.

**Fig. 2:**
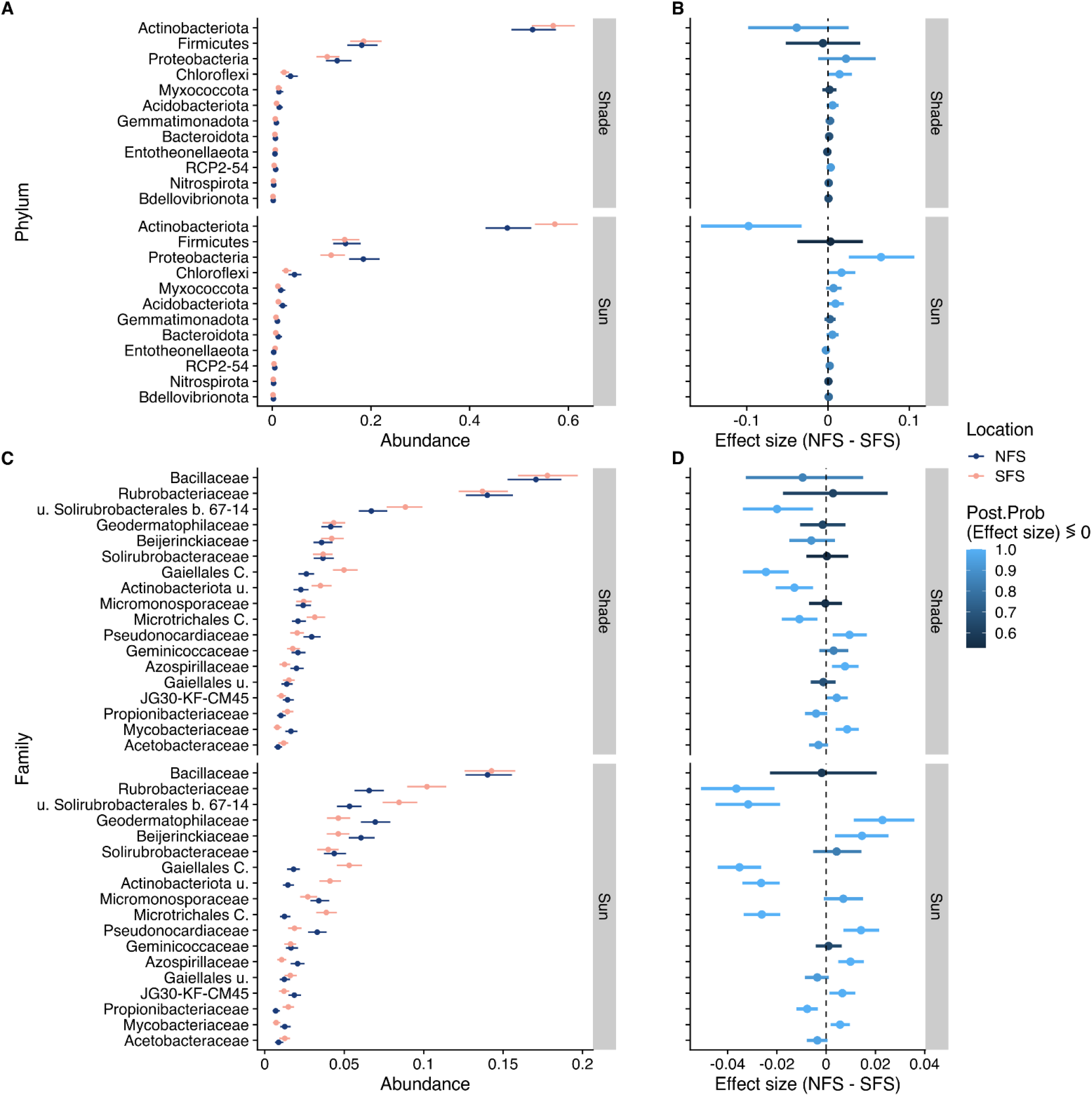
Differential abundance of bacterial taxa in NFS and SFS stations. Relative abundances of bacterial taxa at phylum and family level were analyzed using multilevel modelling. A) Posterior summaries of bacterial phyla from the logistic model [*abundance ∼ phylum * location * light + (1*|*station)*], zero-inflated beta likelihood. B) Differential abundance of phyla obtained from the posterior distributions shown in panel A. C) Posterior summaries of bacterial families from the logistic model [*abundance ∼ family * location * light + (1*|*station)*], zero-inflated beta likelihood. D) Differential abundance of families obtained from the posterior distributions shown in panel C. Points denote the model’s best estimate, and the line indicates a 95% credible interval. Posterior probabilities of the effect size were calculated relative to point null.

### 3.3. Diversity of rhizosphere microbial communities

Analysis of alpha diversity of SFS NFS sun and shade station rhizosphere communities revealed that there was no substantial difference in OTU richness of microbial communities (<u>Fig. 3</u>). The Shannon’s diversity index and inverse Simpson’s index indicate higher diversity at NFS sun stations compared to shade (<u>Fig. 3</u>).

**Fig. 3:**
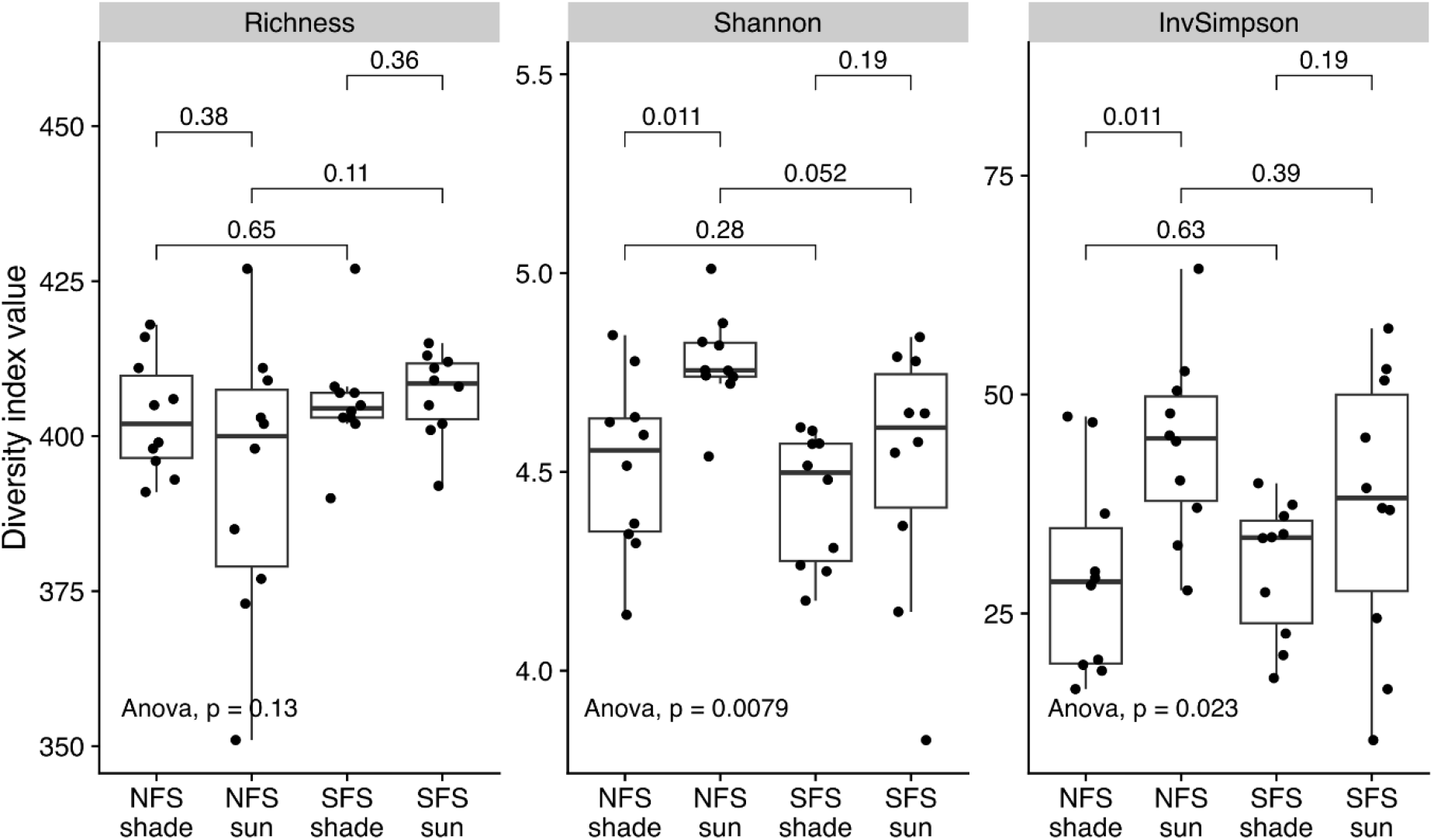
Alpha diversity of rhizosphere microbial communities at SFS NFS sun and shade stations. A) OTU richness. B) Shannon index. C) Inverse Simpson’s index. Low-abundance OTUs and singletons were excluded from the analysis. Points denote original observations. Pairwise comparison p-values are from the Wilcoxon signed-rank test; pairwise p-values were adjusted for multiple comparisons using the ‘holm’ method.

## 4. Discussion

Under the ongoing changes in climate, we can gain knowledge for plant stress adaptation from how the microbially-mediated ecosystem functions at the wild barley rhizosphere from the natural laboratory of EC SFS NFS slopes. Here we studied bacterial distribution on the slopes, sun and shade areas using 16S rRNA analysis of V3-V4 regions. Although the resolution of 16S rRNA sequencing is limited to assignments to the level of genus, it is currently a standard approach in profiling bacterial communities and thus enabled us at least to explore patterns at coarse phylogenetic resolution. While the NFS and SFS shaded areas bacterial distribution didn’t differ, our results show significant differences between the NFS and the SFS sun areas (Fig. 1, 2, and 3). The families *Geodermatophilaceae, Beijerinckiaceae*, and *Pseudonocardiaceae* are dominant in the NFS sun area, and the families *Rubrobacteriaceae*, unclassified *Solirubrobacterales* bacterium 67-14, unclassified *Actinobacteriota*, class *Gaiellales* dominate at the SFS sun area (Fig. 1 A, B, C and D, Fig 2 A and B). Likewise, both Shannon and inverse Simpson’s diversity indices are higher at the NFS sun area compared to the NFS shaded area (Fig. 3 A, B and C).

The results support our former studies on microbial metabolic properties [17], which indicated significantly higher levels of ACC utilization, biofilm formation, phosphorus solubilization and moderately halophilic bacteria content from the SFS slope in comparison to rhizosphere samples from the NFS slope [17]. Yet the current findings also draw attention to the interslope differences of the microbial profiles, which was not the case regarding the five spore-forming taxa, as their properties did not differ between the sun and shaded areas.

Our results are in accordance with former studies at the EC showing the sharp NFS SFS ecological microclimatic divergence, which causes differences in genome size of the carob tree *C. siliqua*, the annual legume *Lotus peregrinus, Oryzaephilus surinamensis* and *Cyclamen persicum* [19]. Similarly, long-term microclimatic stress initiated inter- and intraslope polymorphism in *Nostoc linckia* and differences in GC content in *Bacillus simplex* populations, as well as changes in the content of retrotransposons in the wild barley [19]. The interslope divergence in the whole genome of the small crucifer *Ricotia lunaria* showed upregulation of drought resistance genes on the SFS, and shade resistance and upregulation of photosynthesis genes on the NFS shade. Taken together, the results suggest adaptive patterns to stress conditions. It is clear that the interslope divergence patterns of the EC model, which parallel global patterns of biodiversity and dynamics at the microscale, suggest that these changes are channelled by natural selection, and studies at the contrasting slopes could highlight further effects of global warming at the actively evolving microsites.

The results show that based on the 16S rRNA analysis, there was no substantial difference between diversity indices in SFS sun and shaded areas (Fig. 3 A, B and C). However, it has to be considered that studies of the 16S rRNA variable regions can only show a fraction of the potential variability of the bacterial distribution. It is well known that bacteria in harsh environments are susceptible to modifications, which lead to the emergence of new genetic variance [1]. The modifications can either be short-term adaptations or long-term evolution, and the genetic changes of rhizobacteria can be acquired through various mechanisms including, but not limited to, horizontal gene transfer [21], DNA methylation, and random mutagenesis [22-24] – the changes not reflected in the 16S rRNA gene [1]. More studies have to be performed to unravel the actual inter- and intra-slope differences. However, the results obtained here confirm that new phenotypes can be generated under distinct environmental conditions, which gives the possibility of designing microbial consortia [9], and the EC biomes can be seen as reservoirs of genetic potential that may be mined for identifying microbiome-associated phenotypes. This is the basis of the rationale for designing synthetic communities of microorganisms with wide-ranging, consistent and long-lasting plant growth-promoting traits [1, 25-27]. Hence, the results on the bacterial distribution are an important step towards using transcriptomics, metabolomic profiles and selective plating for figuring out key strains and the supporter strains that strengthen the ecological functions of the key strains, focusing on the PGPR synthetic consortia application to mitigate biotic and abiotic stress factors in future agricultural systems.

## Acknowledgements

Drs. Anastasiia Fetsiukh and David Clapham are gratefully acknowledged for genomic library preparation and critically reading the manuscript. This study was supported by the Swedish Research Council VR 2017-05524 and VR2021-05471 to ST. Sequencing was performed by the SNP&SEQ Technology Platform in Uppsala. The facility is part of the National Genomics Infrastructure (NGI), Sweden, and the Science for Life Laboratory. The SNP&SEQ Platform is supported by the Swedish Research Council and the Knut and Alice Wallenberg Foundation. The computations and data handling were enabled by resources provided by the National Academic Infrastructure for Supercomputing in Sweden (NAISS), partially funded by the Swedish Research Council through grant agreement no. 2022-06725.

## SUPPLEMENTARY INFORMATION

### Mothur batch file commands

set.dir(output=$OUTDIR)

make.file(inputdir=$READS, type=gz, prefix=$PREFIX)

make.contigs(file=current, oligos=./v3v4.oligos, processors=60)

summary.seqs(fasta=current, processors=4)

screen.seqs(fasta=current, count=current, maxambig=0, minlength=402, maxlength=428, maxhomop=8, processors=60)

summary.seqs(fasta=current, count=current, processors=4)

unique.seqs(fasta=current, count=current)

summary.seqs(fasta=current, count=current, processors=4)

align.seqs(fasta=current, reference=./silva.v3v4.align, processors=60)

summary.seqs(fasta=current, count=current, processors=4)

screen.seqs(fasta=current, count=current, start=40, end=17052, processors=60)

filter.seqs(fasta=current, vertical=T, trump=.)

unique.seqs(fasta=current, count=current)

pre.cluster(fasta=current, count=current, diffs=3)

chimera.vsearch(fasta=current, count=current, dereplicate=t)

summary.seqs(fasta=current, count=current, processors=4)

classify.seqs(fasta=current, count=current, reference=./silva.v3v4.align, taxonomy=./silva.nr_v138_1.tax, processors=60)

remove.lineage(fasta=current, count=current, taxonomy=current, taxon=Chloroplast-Mitochondria-unknown-Archaea-Eukaryota)

summary.tax(taxonomy=current, count=current)

rename.file(fasta=current, count=current, taxonomy=current, prefix=$PREFIX.final)

dist.seqs(fasta=current, cutoff=0.03)

cluster(column=current, count=current, cutoff=0.03)

make.shared(list=current, count=current, label=0.03)

classify.otu(list=current, count=current, taxonomy=current, label=0.03)

